# Microbial biomass, composition, and functions are responsible for the differential removal of trace organic chemicals in biofiltration systems

**DOI:** 10.1101/2021.04.22.440850

**Authors:** Lijia Cao, David Wolff, Renato Liguori, Christian Wurzbacher, Arne Wick

**Affiliations:** Chair of Urban Water Systems Engineering, Technical University of Munich, Munich, Germany; Federal Institute of Hydrology (BfG), Koblenz, Germany; Department of Science and Technology, Parthenope University of Naples, Napoli, Italy

**Keywords:** sand filters, metagenome, amplicon sequencing, micropollutants, biotransformation, water treatment, microbiome

## Abstract

Biofiltration processes help to remove trace organic chemicals (TOrCs) both in wastewater and drinking water treatment systems. However, the detailed TOrCs biotransformation mechanisms as well as the underlying drivers behind the variability of site specific transformation processes remain elusive. In this study, we used laboratory batch incubations to investigate the biotransformation of 51 TOrCs in eight bioactive filter materials of different origins treating a range of waters, from wastewater effluents to drinking water. Microscopy, 16S rRNA amplicon and whole metagenome sequencing for assessing associations between the biotransformation rate constants, microbial composition and genetic potential complemented chemical analysis. We observed strong differences in the mean global removal of TOrCs between the individual sand filters (−1.4% to 58%), which were mirrored in overall biomass, microbial community composition, and enzyme encoding genes. From the six investigated biomass markers, ATP turned out to be a major predictor of the mean global biotransformation rate, while compound specific biotransformations were correlated with the microbial community composition. High biomass ecosystems were indicated in our systems by a dominance of Nitrospirae, but individual TOrC biotransformation was statistically connected to rare taxa (< 2%) such as *Hydrogenophaga*, or indiviudal functions such as the enoyl-CoA hydratase/3-hydroxyacyl-CoA dehydrogenase encoding genes. In general, this study provides new insights into so far rarely addressed variability of TOrCs biotransformation. We propose novel biological indicators for the removal performance of TOrCs in biofiltration systems, highlighting the role of living biomass in predicting and normalizing the global transformation, and the role of the microbial community for the individual transformation of TOrCs in engineered and natural systems.

**Contribution to the Field Statement:** Trace organic chemicals (TOrCs) are an emerging problem in the aquatic environment that has attracted global attention over the last decade. Recent research efforts on this topic have increased our knowledge on the transformation of TOrCs and various technologies have been developed to improve their removal. In this study, we investigated a wide range of biotransformation of TOrCs by eight sand filter materials from wastewater and water treatment plants. Biotransformation rate constants were calculated using first-order kinetics to evaluate TOrC removal performance. We reevaluated the role of biomass and could thus explain a greater part of the global TOrC removal performance. The remaining variation in removal rates of individual compounds correlated with the microbiome of the biofilter. Rare biosphere lineages and specific enzyme categories genes were correlated with the removal of certain compounds. In summary, our research identified future indicators for successful biotransformation of TOrCs in biofilter systems.

## Introduction

Trace organic chemicals (TOrCs) such as pharmaceuticals, personal care products and pesticides, have raised emerging concerns regarding their effects on the aquatic environment. These anthropogenic compounds usually enter the wastewater system, and may finally end up in the receiving water bodies, leading to their frequent detection in surface water, ground water and even drinking water at the concentration ranging from few ng·L^-1^ to several μg·L^-1^ (Yang *et al*. 2017; Montiel-León *et al*. 2018; Tröger *et al*. 2018). For example, the widely used artificial sweetener acesulfame has been detected worldwide in wastewater treatment plants (WWTPs) at 10 to 100 μg·L^-1^, and consequently, appears in rivers and groundwater with concentrations up to the double-digit μg·L^-1^ range (Castronovo *et al*. 2017). The effects of residual TOrCs on aquatic ecosystems and the human health have been studied in recent years. For example, Shao *et al*. (2019) found that ten micropollutants with concentrations from 1.1 to 14.4 mg·L^-1^ caused toxicity on zebrafish embryos, aquatic invertebrates and algae. However, there are few examples showing toxic effects also in the sub mg·L^-1^ range for personal care products, hormones, antibiotics, biocides and pesticides (Vieno *et al*. 2014; Yang *et al*. 2017).

In general, WWTPs provide the initial opportunity for removing TOrCs and preventing significant environmental exposure. However, conventional wastewater treatment processes are not originally designed to eliminate diverging TOrCs (Grandclément *et al*. 2017). WWTPs are an important source for the entry of TOrCs; therefore, engineering solutions to improve the removal of TOrCs are needed. Recently, advanced physico-chemical treatment options such as ozonation and activated carbon have been developed and applied at full scale (Kosek *et al*. 2020). In addition, membrane filtration options and advance oxidation processes are also able to efficiently remove TOrCs but are cost intensive (Kanaujiya *et al*. 2019). Conventional activated sludge (CAS) reduces the overall load of micropollutants by both sorption and biodegradation processes but many compounds are only partially removed or persistent (Fischer *et al*. 2014). It has been shown that some of these more persistent compounds are well degraded in subsequent biofiltration systems (Paredes *et al*. 2016; Devault *et al*. 2021). For instance, the removal efficiencies of diclofenac by activated sludge treatment are often below 30% (Zhang *et al*. 2008), while in slow sand filtration systems the removal rate was 40 to almost 80% (Matamoros *et al*. 2007; Casas *et al*. 2015).

Microbial communities bear a high potential to eliminate TOrCs via enzymatic degradation processes, and many recent studies have focused on the influences of various operation parameters to optimize TOrCs removal performance. So far, especially redox conditions (Oberleitner *et al*. 2020), carbon and nitrogen availability (Moe *et al*. 2001; Zhang *et al*. 2019), hydraulic retention time (Priya *et al*. 2015), and the filter material (Paredes *et al*. 2016) have been reported to affect the efficiency of different biofiltration systems. These studies have contributed to our understanding of micropollutant biotransformation in engineered systems. However, all of the aforementioned operating conditions did not have a direct impact, they in fact affect the microbial communities and thereby the diversity, abundance and function of microorganisms.

To date, the micropollutant biotransformation mechanisms as well as the underlying drivers behind the degradation processes remains elusive. Therefore, to uncover the microbial “black box”, additional studies regarding the associations between the abundance of taxa, enzymes, pathways and TOrCs biotransformation are required. Johnson *et al*. (2015) for example, investigated the taxonomic biodiversity and biotransformation rates of ten micropollutants in ten full-scale WWTPs, and found biodiversity was positively associated with the collective removal rates of TOrCs. To explore the biodiversity in more detail, Wolff *et al*. (2018) divided the general microbial community into a core and a specialized community based on defined filter criteria and statistical selection, comparing their composition with micropollutant removal in five different biological wastewater treatment systems. The results demonstrated the significant correlation between the relative abundance of specialized community members and the removal rates of certain compounds. For example, the abundances of the genera *Luteimonas*, *Roseibaca* and *Phenylobacterium* might be indicative for the degradation of metoprolol, 10,11-dihydro-10-hydrocarbamazepine and diclofenac under aerobic condition. Functional enzymes and degradation pathways are often studied in pure culture or individual compound biodegradation studies (Kjeldal *et al*. 2019), as the enzymatic reactions involving diverse microorganisms are quite intricate, making it hard to interpret the degradation pathways (Achermann *et al*. 2018; Zumstein *et al*. 2019). Despite these initial studies, the biotransformation mechanisms of various TOrCs in the biological filtration system on the microbial community level are rarely addressed so far, only few studies focused on their removal efficiencies and influencing factors (Zearley *et al*. 2012; Rattier *et al*. 2014; Carpenter *et al*. 2017).

In this study, biotransformation performance of eight different biological active sand filter materials from wastewater and water treatment plants for transforming 51 polar TOrCs were investigated experimentally. We coupled biotransformation rates with multiple microbial parameters, which involved the detailed analysis of degrading microorganisms, their functional genes and transformation pathways. The aim of this study is: (1) to assess and compare the transformation efficiencies of different types of biological active systems in a fully controlled laboratory setup, (2) to evaluate the influence of the microbial community composition on the biotransformations, (3) to suggest novel biological indicators for a more comprehensive evaluation of individual or global TOrCs removal efficiencies. For this purpose, we put the eight sand filters under strictly controlled laboratory conditions, thereby excluding any influence of the above-mentioned parameters (redox conditions, carbon and nitrogen availability, hydraulic retention time) on differences in biotransformation rates. Hence, we focused on the transformation potential of the microbial communities at identical conditions, which allowed us to directly link biotransformation rates with taxa, functional genes and biomass markers. We hypothesize that differences in biotransformation rates will be mirrored by the a) microbial community, and their b) enzymatic repertoire encoded in the metagenome. Moreover, we evaluated which role biomass plays in the normalization of global transformation performance. This may lead to the identification of biological parameters for the removal of TOrCs, individually or globally.

## Materials and methods

### Trace organic compounds

51 polar TOrCs which have been typically found in municipal wastewater were selected as target compounds in this study (Table 1), including pesticides, pharmaceuticals, and personal care products with different biodegradabilities in CAS treatment. For example, acyclovir, atenolol, caffeine, and ibuprofen can be relatively easily degraded (Prasse *et al*. 2011; Ferrando-Climent *et al*. 2012; Xu *et al*. 2017a; Chtourou *et al*. 2018), while carbamazepine and diclofenac are persistent and difficult to be biotransformed (Kruglova *et al*. 2014; Zhang *et al*. 2008), and some compounds such as emtricitabine and cetirizin have only been rarely studied so far. All TOrCs reference standards used were of analytical grade (Sigma-Aldrich) with a minimal purity of 98%.

**Table 1.** Names, usage and abbreviations of the 51 TOrCs analyzed in this study and the statistical differences of their *k_biol_* after normalization with ATP.

| Compound | Usage | Abbreviation | Kruskal-Wallis-Test<br>(adjusted $p$-value) | | | |
| --- | --- | --- | --- | --- | --- | --- |
|  |  |  | normalizaed |  | non-normalized |  |
|  |  |  | between<br>7 sands | HBG vs<br>LBG | between<br>7 sands | HBG vs<br>LBG |
| 10-hydroxycarbamazepine | Antiepileptics | 10-OH-CBZ | * | ** | n.s. | * |
| 2-hydroxycarbamazepine | Antiepileptics | 2-OH-CBZ | n.s. | n.s. | ** | * |
| 3-hydroxycarbamazepine | Antiepileptics | 3-OH-CBZ | * | n.s. | * | * |
| Acesulfame | Sweetener | ACE | n.s. | n.s. | * | n.s. |
| Acridone | TP of 10-OH-CBZ | - | - | - | - | - |
| Acyclovir | Antivirals | ACV | n.s. | * | ** | * |
| Amisulprid | Neuroleptics | - | n.s. | n.s. | ** | * |
| Atenolol | Beta-Blockers | ATN | * | ** | n.s. | * |
| Azithromycin | Antibiotics (macrolides) | AZI | * | ** | n.s. | * |
| Benzotriazole | Industrial | BTA | n.s. | n.s. | * | * |
| Bezafibrate | Lipid modifying agent | BEZ | n.s. | n.s. | ** | * |
| Caffeine | Psychoactive drug | CAF | - | - | - | - |
| Carbendazim | Fungicide | - | n.s. | n.s. | ** | * |
| Carboxy-Acyclovir | TP of acyclovir | - | * | * | ** | * |
| Carbamazepine | Antiepileptics | CBZ | n.s. | n.s. | * | n.s. |
| Ceterizin | Antihistamine | CTZ | n.s. | n.s. | ** | * |
| Clarithromycin | Antibacterials | CLR | * | ** | n.s. | * |
| Climbazole | Antifungal | CZ | * | n.s. | ** | * |
| Clopidogrel acid | Metabolite of clopidogrel | - | * | n.s. | ** | * |
| Clopidogrel | Antiaggregant (pro-drug) | CLO | * | n.s. | ** | * |
| N,N-Diethyl-meta-toluamide | Insecticide/Repellent | DEET | n.s. | n.s. | ** | * |
| Diclofenac | Antiinflammatory | DCF | * | n.s. | * | * |
| Diuron | Herbicide/Algicide | DIU | * | * | ** | * |
| Emtricitabine | Virostatic agent | EMT | * | * | ** | * |
| Fexofenadine | Antihistamine | FEX | * | * | n.s. | * |
| Flecainide | Antiarrhythmics | FLEC | n.s. | n.s. | ** | * |
| Fluconazole | Antimycotics | FCZ | n.s. | n.s. | * | n.s. |
| Furosemide | Loop diuretics | FRM | * | ** | ** | * |
| Gabapentin | Anticonvulsant drug | GAP | n.s. | n.s. | ** | * |
| Hydrochlorothiazide | Diuretics | HCT | n.s. | n.s. | * | n.s. |
| Ibuprofen | Antirheumatic drug | IBU | - | - | - | - |
| Iopromide | Contrast media | IOP | * | n.s. | * | * |
| Lamotrigine | Anticonvulsant drug | LTG | n.s. | * | * | n.s. |
| Levetiracetam acid | TP of levetiracetam | - | - | - | - | - |
| Lidocaine | Local anaesthetic | LD | n.s. | n.s. | * | * |
| Mecoprop | Herbicide | MCPP | * | ** | ** | * |
| Metoprolol | Beta blockers | MET | n.s. | ** | n.s. | * |
| Oxypurinol | Metabolite of allopurinol<br>(antigout agent) | OP | * | n.s. | n.s. | * |
| Pregabalin | Anticonvulsant drug | PGB | n.s. | * | ** | * |
| Ramiprilat | Antihypertensives | - | - | - | - | - |
| Sitagliptine | Antidiabetics | SG | n.s. | n.s. | ** | * |
| Sulfamethoxazole | Antibacterials | SMX | * | n.s. | n.s. | * |
| Sulpirid | Neuroleptics | SP | n.s. | n.s. | ** | * |
| Terbutylazine | Herbicide | TBA | n.s. | n.s. | ** | * |
| Terbutryn | Herbicide/Algicide | - | n.s. | n.s. | ** | * |
| Torasemid | Diuretics | - | n.s. | n.s. | ** | * |
| Tramadol | Analgesic | TRA | n.s. | n.s. | * | * |
| Trimethoprim | Antibacterials | TMP | * | ** | n.s. | * |
| Valsartan | Antihypertensives | VAL | - | - | - | - |
| Venlafaxine | Psychoanaleptics | VEN | n.s. | n.s. | n.s. | n.s. |
| Xipamide | Diuretics | XPD | n.s. | n.s. | * | n.s. |

TP: transformation products
n.s.: $p \geq 0.05$ ; \*: $0.01 \leq p < 0.05$ ; \*\*: $0.001 \leq p < 0.01$

### Experimental setup and operation

Six different sand materials from rapid sand filters of municipal WWTPs and two materials from slow sand filters used for water treatment were sampled and then stored at 4 °C for approximately three months. The batch experiments were performed in triplicates with all eight sand filters under the same conditions: 20 g of sand material was added to each 250 mL bottle and 80 mL of treated wastewater from the WWTP Koblenz. All batches were aerated with 7 mg·L^-1^ oxygen and continuously shaken at 22 °C and the speed of 100 rpm in the dark. To acclimate the microorganisms, all batches were equilibrated to the experimental conditions for approximately 24 h before the experiment started. The experiment was started by spiking the bottles with a mixture of 51 TOrCs at the initial concentration of 0.5 μg·L^-1^ for each compound, which was within the range of TOrCs concentration in actual wastewater (Luo *et al*. 2014).

### Sampling

Over 72 h incubation, water samples were withdrawn from each bottle at regular intervals resulting in eight time points (0, 1, 4, 8, 12, 24, 48, 72 h) for *k_biol_* calculation (see below). Samples were immediately filtered using 0.45 μm regenerated cellulose membrane (Macherey-Nagel, Germany) and stored at 4 °C for maximum 2 weeks until liquid chromatography tandem mass spectrometry (LC-MS/MS) analysis. At the end of the experiment, the remaining water was removed and the sand materials were collected and stored at −80 °C for DNA extraction. In addition, each sand material was fixed 1:1 with ethanol for fluorescence *in situ* hybridization (FISH) analysis.

### Biomass measurement and FISH analysis

In contrast to the DNA based analysis and the monitoring of the compound removal (see below), the biomass measurements and the cell counting was performed as one measurement per filter material (n = 8). Loss on ignition was determined by drying 15 g of sand filter material at 105 °C overnight, and then placing in a muffle furnace at 440 °C for 4 h, weighing the samples at each step. ATP from 1 g sand filter material was measured by the single tube luminometer Sirius FB12 (Titertek Berthold, Germany) using the ATP Biomass Kit HS (BioThema, Sweden) according to the manufacturer’s instructions with eight measurement points prior and after ATP standard addition. The sand samples were sent to Vermicon AG (Munich, Germany) for cell count measurement and FISH analysis. FISH counting was performed with the sand samples using fluorescence microscopy for following groups: Alphaproteobacteria, Betaproteobacteria, Gammaproteobacteria, Deltaproteobacteria, Epsilonproteobacteria, Actinobacteria, Firmicutes, Cytophaga-Flexibacer-Subphylum, Planctomycetes, Chloroflexi, Nitrospirae, TM7, Archaea. Probe sequences and hybridization conditions were taken from Daims et al., (2006). FISH protocol was performed following Snaidr et al., (1997) with following modifications for sample preparation: 10 g of sand samples were thoroughly vortexed for 5 min. After settlement of the samples at room temperature 200 μl of supernatant were transferred into a 2 ml vial and mixed with 1 ml of PBS buffer. Samples were vortexed, followed by centrifugation at 5000 rpm for 1 min. Supernatant was extracted and transferred into new 2 ml vial, followed by centrifugation at 5000 rpm for 5 min. Supernatant was discarded and remaining pellet was solved in 200 μl of 50% EtOH. At least ten fields of view were counted per measurement using a Zeiss Axioscop 2 epifluorescence microscope equipped with fluorescence filter sets for the dyes FAM, Cy3 and 4,6-Diamidino-2-phenylindol-dihydrochloride (DAPI). More information can be obtained by request from Vermicon AG (Munich, Germany).

### LC-MS/MS measurement and calculation of biotransformation rates

To determine the concentrations of the 51 spiked TOrCs, all filtered samples of the eight time points were used for LC-MS/MS analysis via an Agilent 1260-LC coupled to a SCIEX QTrap 5500-MS according to the method previously described by Falås *et al*. (2016). Along the sampled 72h, biotransformation rate constants were estimated assuming first-order kinetics and negligible sorption according to equation (1), whereby *S* is the TOrCs concentration (μg·L^-1^), *X_Biomass_* is the biomass concentration represented by the ATP (pmol·g^-1^), and *k_biol_* is the normalized pseudo-first-order rate constant in g·(pmol·d)^-1^.

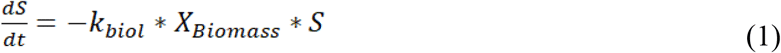

### DNA extraction and sequencing

The DNA of the microorganisms was extracted from the sand filter material, and was used to analyze the present taxa by 16S rRNA amplicon sequencing, and by whole genome shotgun sequencing. Approximately 1.5 g of the frozen filter material from the end of the experiment (24 samples) was used for DNA extraction using the FastDNA SPIN Kit for soil (MP Biomedicals, Eschwege, Germany) according to the manufacturer’s instructions. Concentrations and purity of the individual DNA extracts were measured by microspectrophotometry (NanoPhotometer P330, Implen). 16S rRNA gene amplification, library preparation and sequencing was performed by IMGM Laboratories GmbH (Planegg, Germany). In short, primer pair 341F (CCTACGGGNGGCWGCAG) and 805R (GACTACHVGGGTATCTAATCC) for V3-V4 hypervariable region was used for amplification of the 16S gene. Next, the purified PCR products were normalized to equimolar concentration with the SequalPrep Normalization Plate Kit (Thermo Fisher Scientific) and the DNA library was sequenced on the Illumina MiSeq next generation sequencing system (Illumina Inc.) in paired end mode with 2 x 250 bp. Whole metagenome sequencing was performed by IMGM Laboratories GmbH (Planegg, Germany). In brief, the extracted DNA samples were checked for quality and quantity by a 1% agarose gel (Midori Green-stained) electrophoresis and a Qubit dsDNA HS Assay Kit (Invitrogen). Prior to library preparation, the high molecular weight gDNA was sheared to target fragment size of 550 bp using a Covaris M220 Focused-ultrasonicator (Brighton, United Kingdom). In the following, DNA was prepared for sequencing using the NEBnext UltraTM II DNA library preparation kit for Illumina (New England Biolabs). Sequence data were generated on the Illumina HiSeq 2500 next generation sequencing system with the 2 x 250 bp paired end mode. 16S and whole metagenome sequence data are freely available at ENA under the accession number PRJEB43767.

### Data processing

All 16S rRNA gene amplicons data were processed in R (v 3.6.0) (R Core Team 2016) using the DADA2 (v 1.9.0) pipeline (Callahan *et al*. 2016). Quality trimming, denoising, error-correction, paired-end read merging, chimera removal, and dereplication steps were performed according to the default setting (Table S1). The amplicon sequence variants (ASVs) were taxonomically classified with a naïve Bayesian classifier using the SILVA training dataset (v137) (Quast *et al*. 2013). The ASV and taxonomy tables, along with associated sample metadata were imported into phyloseq (v 1.22.3) (McMurdie and Holmes 2013) for community analysis. Whole metagenome sequencing data was processed using Trimmomatic (v 0.39) (Bolger *et al*. 2014), and the quality of the reads was checked with FastQC (v 0.11.8) (Andrews 2010). The metagenome data was used for a) validating the amplicon results with METAXA2 software (v 2.1.1) (Bengtsson-Palme *et al*. 2015); b) calculating the metagenomic distances between the samples by making a gene-pool comparison/relatedness determination with the k-mer based weighted inner product, kWIP (Murray *et al*. 2017); c) metagenome assembly and functional analysis. For these latter two analysis, we used MEGAHIT (v 1.2.9) (Li *et al*. 2015) to assemble the data, and QUAST (v 5.0.2) (Gurevich *et al*. 2013) to evaluate the assembly quality. For the subsequent functional analysis, SUPER-FOCUS (v 0.31) (Silva *et al*. 2015) was applied using the aligner DIAMOND (v 0.9.34) to obtain the enzyme commission (EC) catogories (Buchfink *et al*. 2015). In addition, taxonomic informed pathways classifications by sample groups were determined by HUMAnN2 (v 2.8.1) (Franzosa *et al*. 2018). Metagenome-assembled genomes (MAGs) were obtained by MetaBAT2 (v 2.15) (Kang *et al*. 2019) and manually refined after assessing the completeness (> 70%) and contamination (< 10%) by CheckM (v 1.0.13) (Parks *et al*. 2015). Taxonomic classification was conducted by GTDB-Tk (Chaumeil *et al*. 2020). Prodigal (v 2.6.3) (Hyatt *et al*. 2010) was used for open reading frames (ORFs) prediction. KofamKOALA (https://www.genome.jp/tools/kofamkoala/) (Aramaki *et al*. 2020) was used to obtain KO annotations for genes predicted by Prodigal. Xenobiotics metabolism was performed by the “Reconstruct Pathway” tool in KEGG mapper (https://www.genome.jp/kegg/mapper.html, accessed February 2021).

### Statistical analysis

All statistical analysis were performed in R environment (v 3.6.0). *cor.test* function was used to test the correlation coefficient *r* between the TOrCs biotransformation performance and different biomass indicators based on the Pearson method. Linear regression was used to model the global compound removal by ATP concentration. Residuals were tested for normality (Shapiro-Wilk normality test), and the distribution was inspected through QQ (quantile-quantile) plot (Fig. S1). A paired two-sample Mann-Whitney-Wilcoxon test (non-parametric) was used to identify significant differences between *k_biol_* values of two sand groups. Kruskal-Wallis test was used to evaluate the differences of individual compound *k_biol_* among all sand samples and between categories. The relationships between DADA2 and METAXA2 data were examined by Mantel tests. The microbial diversity indices were analyzed using the vegan package (v 2.5-6) (Oksanen *et al*. 2019). The species richness was determined by rarefying the amplicon dataset to the smallest sample (5799 reads) through the “rrarefy” function (Fig. S2). Chao1 and Inverse Simpson index (performed on full dataset prior to rarefaction) were used to present community richness and alpha diversity, respectively. Community compositions were compared using Bray-Curtis dissimilarities on ASV abundances and presented using NMDS ordinations. Ordinations and heatmaps were done in the R package “ampvis2” (v 2.4.6) (Andersen *et al*. 2018). The analysis of correlations between TOrCs biotransformation rates and microorganisms or functional genes was based on Pearson correlations outlined in the Rhea script collection (Lagkouvardos *et al*. 2017). Before doing the correlation analysis, we tested the differences between each compound *k_biol_* and zero (criterion 1), and set the minimum removal percentage to 10% (criterion 2). As a consequence, the *k_biol_* of carbamazepine, fluconazole and xipamide were excluded from further analysis, since either criterion 1 or 2 was not met in any of the batches. Differential functional genes and pathway analyses were conducted by DESeq2 (v 1.29.5) (Love *et al*. 2014).

## Results

### Correlation of TOrCs biotransformation performance with biomass

Physical, chemical and biological characteristics of eight sand filters are shown in Table 2. Materials of these filter included quartz gravel, quartz sand, anthracite and pumice. Loss on ignition, ATP, DNA concentration and cell counts served as biomass indicators. Biomass was high in Friedrichshafen, Eriskirchen and Wangen (> 25 pmol/g ATP, > 40 mg/g Loss on ignition), whereas the remaining five biofilters were low in biomass, with the lowest measurements in the filter materials from drinking water systems (BWA and IFW; < 1.5 pmol/g ATP, < 9 mg/g Loss on ignition) (Table 2, Fig. 1b). The biotransformation rate constants of TOrCs were calculated excluding six compounds (i.e., acridone, caffeine, ibuprofen, levetiracetam acid, ramiprilat and valsartan) which did not fit the first-order kinetics. Moreover, carbamazepine, fluconazole, and xipamide had non-significant *k_biol_* and showed less than 10% removal in all sand samples. Wangen achieved the best overall TOrC parent compound removal with a global mean removal percentage of 58 ± 0.64%, followed by Friedrichshafen (49 ± 3.4%%) and Eriskirchen (45 ± 1.2%) (Fig. 1a). Hungen and Stuttgart showed similar global removal percentages of 28% and 26%, followed by Moos with 14%, respectively. In the drinking water filters IFW and BWA, there were almost no TOrCs degraded (removal below 0.5%). These differences in global transformation potential were highly correlated with biomass expressed as ATP concentration (Pearson *r* = 0.92, *p* < 0.001), followed by living cell counts (Pearson *r* = 0.76, *p* < 0.001), while loss on ignition, which has been traditionally used for the *k_biol_* normalization showed no correlation (Table 2). The mean global *k_biol_* could be predicted by ATP concentration (logarithmic: ln(ATP), LM, *t* = 8.3, adjusted R^2^ = 0.76, *p* < 0.001; or linear: LM, *t* = 10.9, adjusted R^2^ = 0.84, *p* < 0.001, Fig. 1c). The linear model showed a better fit, but the distribution of residuals was non-normal (Fig. S1c). In addition to *k_biol_*, the overall average removal percentage showed a clear relationship with ATP concentration, this time on a natural logarithmic scale (LM, *t* = 13.9, adjusted R^2^ = 0.90, *p* < 0.001, Fig. 1d). Therefore, we used ATP for *X_Biomass_* normalization (equation 1) of *k_biol_* for the clustering analysis and the following correlation analysis. According to the ATP biomass, the samples with high biomass (Eriskirchen, Friedrichshafen and Wangen, named as HBG) had a high general biotransformation ability, while the biotransformation potential of the samples with low biomass (Hungen, Moos and Stuttgart, named as LBG) was relatively low, which can be also seen across individual compounds (Fig. 2a). After ATP normalization, however, the individual differences between filter materials and compounds became less pronounced (Fig. 2b, except IFW). When comparing DW with the other two groups, their *k_biol_* (mean value of individual substances in each group) remained different (Mann-Whitney-Wilcoxon test, W = 1305, *p* = 0.02 for HBG; W = 1318, *p* = 0.01 for LBG). However, when testing for differences in biotransformation of LBG and HBG, there was no significant difference in the mean global *k_biol_* between the two groups (Mann-Whitney-Wilcoxon test, W = 1305, *p* = 0.33). This indicated their mean biotransformation performance were comparable per biomass unit, independent of other parameters such as microbial community composition. Differences between the *k_biol_* of individual compounds persisted when comparing seven sand samples (IFW was excluded: the values close or below zero caused superimposed signals when normalized, Fig. 2b) or HBG and LBG. Overall, 19 compounds showed significant differences in seven sand samples, and 15 substances showed significantly differential *k_biol_* between HBG and LBG (Kruskal-Wallis test, adjusted *p* < 0.05) (Table 1). For comparison, we found 35 and 38 substances to differ in their *k_biol_* before normalization for the sand filter materials and HBG vs. LBG, respectively.

**Figure 1.**
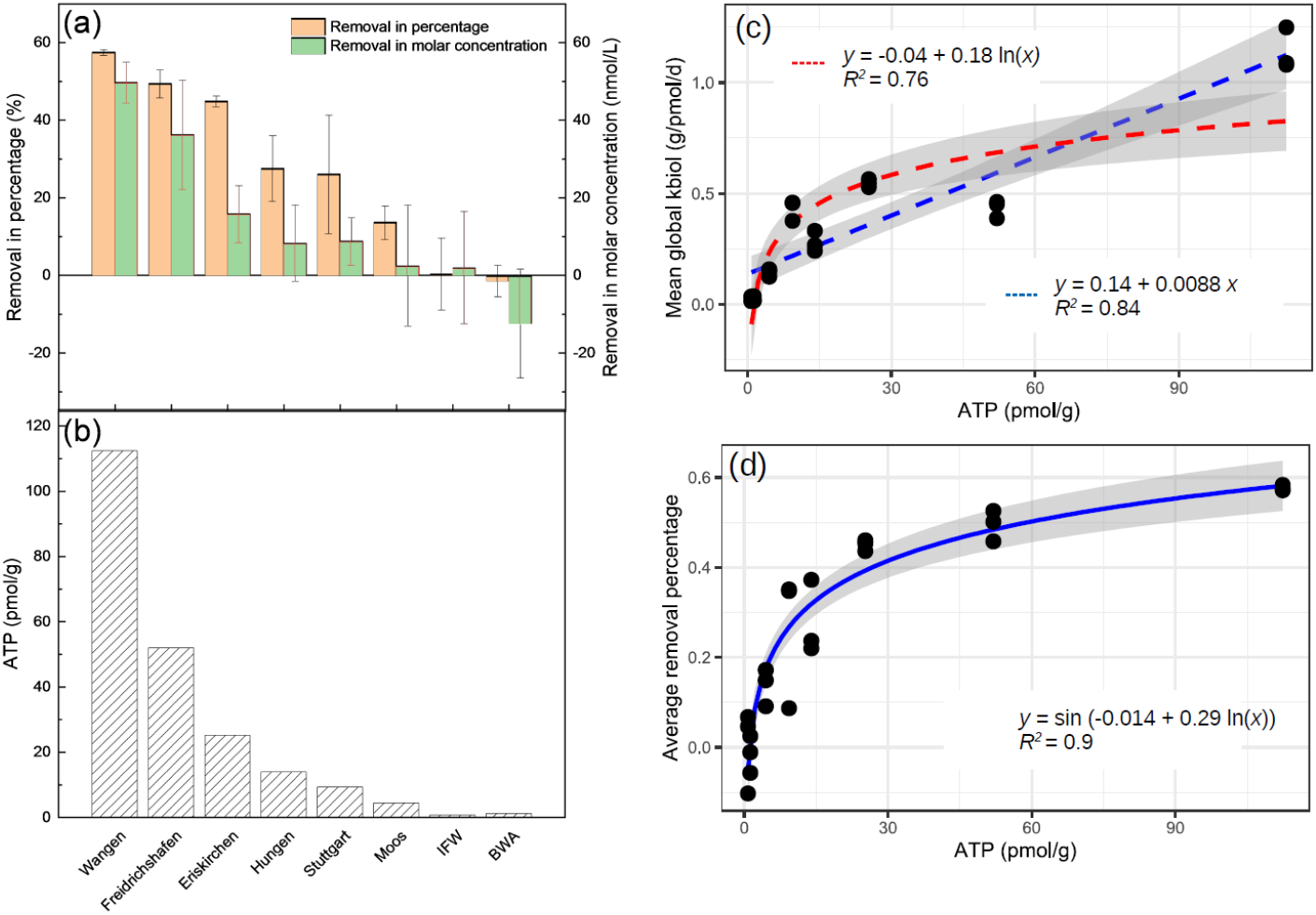
(a) Average global removal of 51 TOrCs by eight sand filter materials in percentage and molar concentration; (b) ATP concentration in eight sand filter samples; Regression (blue/red line) for predicting (c) mean global biotransformation rate constants (*k_biol_*) and (d) average removal percentage of TOrCs from ATP concentration, gray shadow represents the 95% confidence interval.

**Figure 2.**
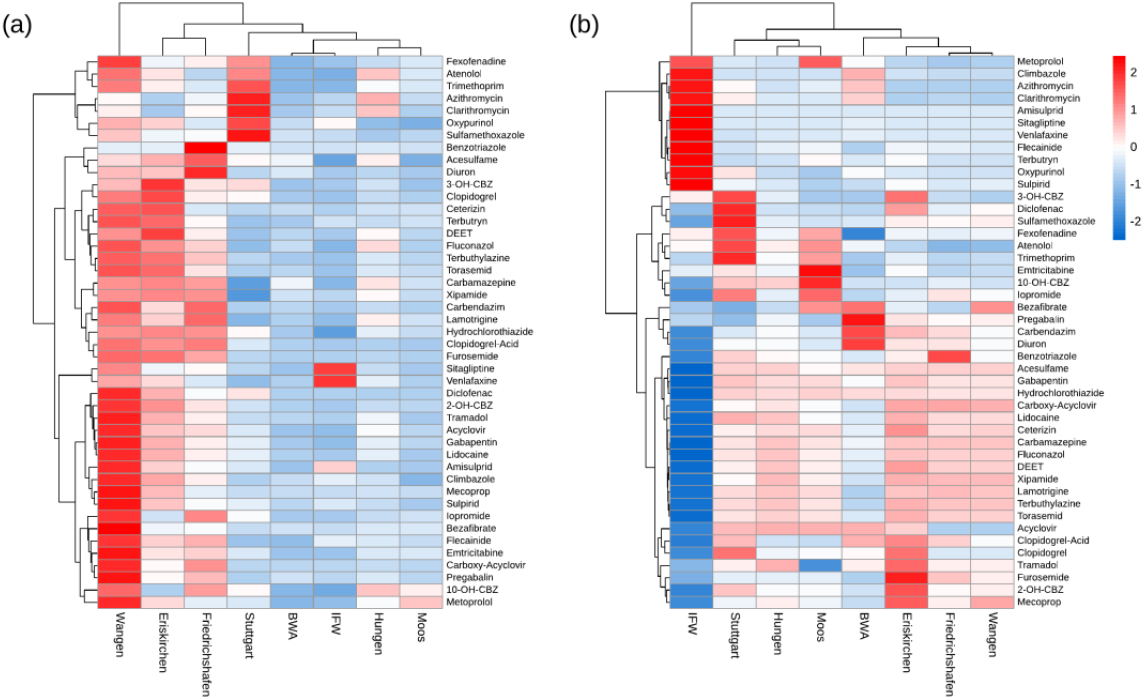
(a) Non-normalized and (b) normalized biotransformation rate constants (*k_biol_*) of 45 TOrCs by eight sand filter materials. The low ATP concentration in the IFW sand (0.76 pmol/g) made the normalized results appear superimposed. The *k_biol_* values were scaled within appropriate range (−2, 2) for better visualization. Clustering used the Manhattan distance metric and Ward’s minimum variance method (Ward.D2). Six TOrCs were excluded as they did not fit the first-order kinetics.

**Table 2.** Biological characterization of eight sand filter materials and correlation test between TOrCs removal and biomass indicators. The significance of the Pearson’s *r* correlation coefficient was adjusted for multiple comparisons by the bonferroni method.

| Sampling site | Sand filter type | Material | Loss on ignition (mg biomass/g) | ATP (pmol/g) | DNA concentration (ng/g) | Viable cell counts (cells/g) | Dead cell counts (cells/g) | Total cell counts (cells/g) | Alpha diversity |  |
| --- | --- | --- | --- | --- | --- | --- | --- | --- | --- | --- |
|  |  |  |  |  |  |  |  |  | Chao1 | Inverse Simpson |
| Hungen | No addition | Quartz gravel | 2.23 | 13.98 | 1.6 | 1.58E+08 | 8.10E+07 | 2.39E+08 | 2197.90 | 105.07 |
| Stuttgart | No addition | Anthracite | 13.61 | 9.34 | 3.6 | 9.81E+08 | 3.79E+08 | 1.36E+09 | 993.67 | 120.87 |
| IFW | Artificial groundwater recharge | Quartz sand | 3.40 | 0.76 | 1.3 | 6.27E+07 | 1.46E+08 | 2.09E+08 | 824.76 | 10.59 |
| BWA | Waterworks | Quartz sand | 9.02 | 1.25 | 1.9 | 1.60E+08 | 1.00E+08 | 2.60E+08 | 787.93 | 8.01 |
| Moos | Precipitant addition | Quartz gravel | 2.96 | 4.48 | 1.2 | 2.35E+08 | 1.34E+08 | 3.69E+08 | 826.78 | 48.31 |
| Friedrichshafen | Precipitant and carbon source addition | Anthracite | 314.19 | 52.00 | 3.9 | 1.57E+09 | 3.30E+08 | 1.90E+09 | 2072.95 | 290.81 |
| Eriskirchen | Precipitant addition | Anthracite | 179.46 | 25.27 | 3 | 5.69E+08 | 2.23E+08 | 7.92E+08 | 1357.69 | 65.54 |
| Wangen | Precipitant addition | Pumice | 41.15 | 112.41 | 3.2 | 1.34E+09 | 2.90E+08 | 1.63E+09 | 1069.59 | 38.19 |
| Non-normalized biotransformation rate constants ($k_{biol}$) | correlation coefficient | | 0.25 | 0.92 | 0.64 | 0.76 | 0.61 | 0.75 | 0.16 | 0.15 |
| | adjusted $p$-value | | 1.00 | < 0.001 | 0.006 | < 0.001 | 0.01 | < 0.001 | 1.00 | 1.00 |

### Microbial community composition of sand filter biofilms

We investigated the microbial community composition of all incubations using 16S rRNA amplicon sequencing and metagenomics. For the amplicon data, all sequences were clustered into a total of 23147 bacterial ASVs after the filtering step, showing obvious differences among the sand filters at the phylum level (Fig. 3a). Apart from a high abundance of Proteobacteria in all filter materials, Nitrospirae dominated in the HBG with mean read abundances of 27.6%, followed by Bacteroidetes (11.5%). In the LBG, Nitrospirae only accounted for 0.7%. Microbial community composition was further cross-validated by microscopy (FISH) and metagenome classifications (METAXA2) (Fig. 3a). In particular, the community composition matrix that resulted from the metagenome (METAXA2) was highly correlated with the composition matrix generated (at much deeper resolution) with the ASV amplicon data (Mantel tests, *r* = 0.96, *p* = 0.001). The FISH results confirmed that the relative proportion of cells and their respective biomass (assuming the same cell size) follow the results of the DNA based data (with exceptions; e.g., for the Nitrospirae community in Stuttgart).

**Figure 3.**
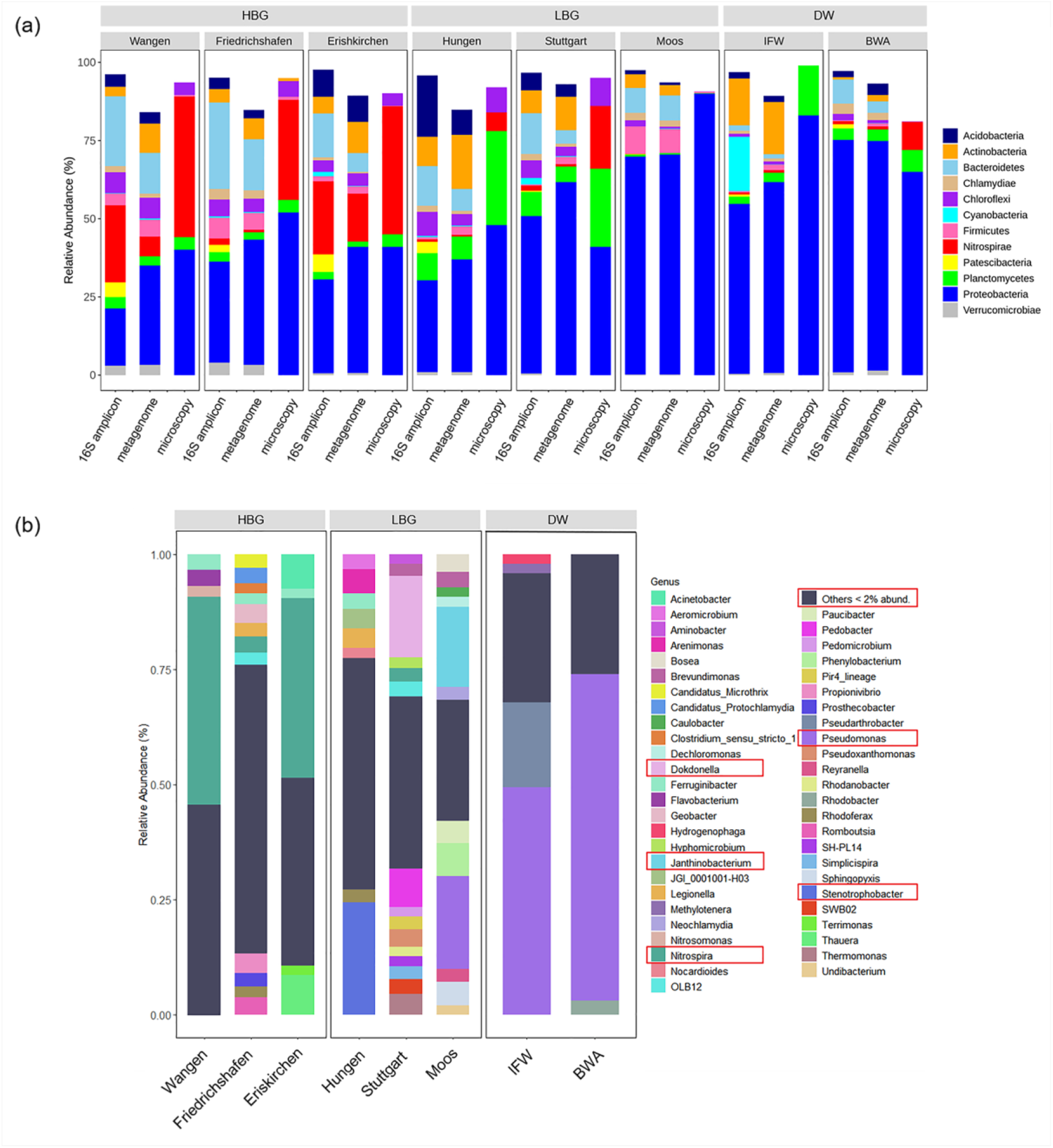
Taxonomic composition of eight sand filter materials. (a) Comparison of taxa from microscopy (FISH) vs. 16S amplicon (DADA2) vs. metagenome (METAXA2); (b) Relative abundance of microorganisms at the genus level.

At the genus level, a high degree of variation in the taxonomic composition between the sand samples was observed (Fig. 3b, Fig. S3). In the HBG, there was a high amount of *Nitrospira* in Wangen and Eriskirchen with mean relative abundances of 45.2% and 39.0%, respectively. In Friedrichshafen (which was operated for post-denitrification), *Nitrospira* only accounted for 3.6% and other lineages became more abundant (e.g., *Propionivibrio*, *Geobacter*, *Romboutsia*, *Legionella*). In LBG, the three sand samples exhibited largely different microbial composition without sharing abundant genera. *Stenotrophobacter* (24.4%) was prominent in Hungen and *Dokdonella* (17.8%) dominated the Stuttgart filter materials. *Pseudomonas* (20.1%) and *Janthinobacterium* (17.3%) were dominant in Moos. In contrast to HBG and LBG, DW showed a stark dominance of *Pseudomonas* with a mean relative abundance of 60.1%, which also resulted in the lowest biodiversity indices (Table 2). In a multivariate analysis of the community composition (presented as ASV or as k-mers derived from the metagenome) we could recover a separation into HBG and LBG (Fig. S4, adonis: for ASV, R^2^ = 0.21, *p* = 0.001; for metagenomics k-mers (kWIP), R^2^ = 0.19, *p* = 0.001).

### Correlation of TOrCs biotransformation with microorganisms and functional genes

Finally, we confirmed that the community composition may be linked to the biotransformations observed in the systems using a Mantel test for the ASV matrix and the (normalized) *k_biol_* values of the incubations (*r* = 0.50, *p* = 0.001). Subsequently, we were interested if there are any specific linkages between microbes and biotransformations. For this, Pearson’s coefficients *r* were used to find hypothetical linkages between the *k_biol_* and relative abundances of a) microbial genera (Fig. 4a) or b) functional genes (Fig. 4b, see paragraph below). In total, there were 62 genera (as a data reduction step, all ASV were collapsed to genera for this purpose) that showed either a significantly positive or negative correlation with the biotransformation rates of TOrCs (with a cutoff of abs(*r*) > 0.7, adjusted *p* < 0.05, observation > 9) (Fig. 4a). All of these (except *Pseudomonas*) ranked below 2% relative abundance and can be regarded as rare taxa. Thirty genera showed highly positive correlations with more than one TOrC, such as *Denitratisoma, Hydrogenophaga* and *Ideonella*. Four genera correlated with single TOrCs, i.e., *Litorilinea, Novosphingobium, Paludibaculum, Phaeodactylibacter* were only positively associated with the biotransformation of diclofenac, acyclovir, sulfamethoxazole and bezafibrate, respectively. *Nitrospira*, the dominant genus in the HBG, had no significant correlation with any compound (only when non-normalized *k_biol_* values are considered we found multiple negative correlations). Examples for the negative correlation were *Conexibacter, Ferruginibacter, Intestinibacter, Methylibium, Pseudomonas* and *Terrimonas*.

**Figure 4.**
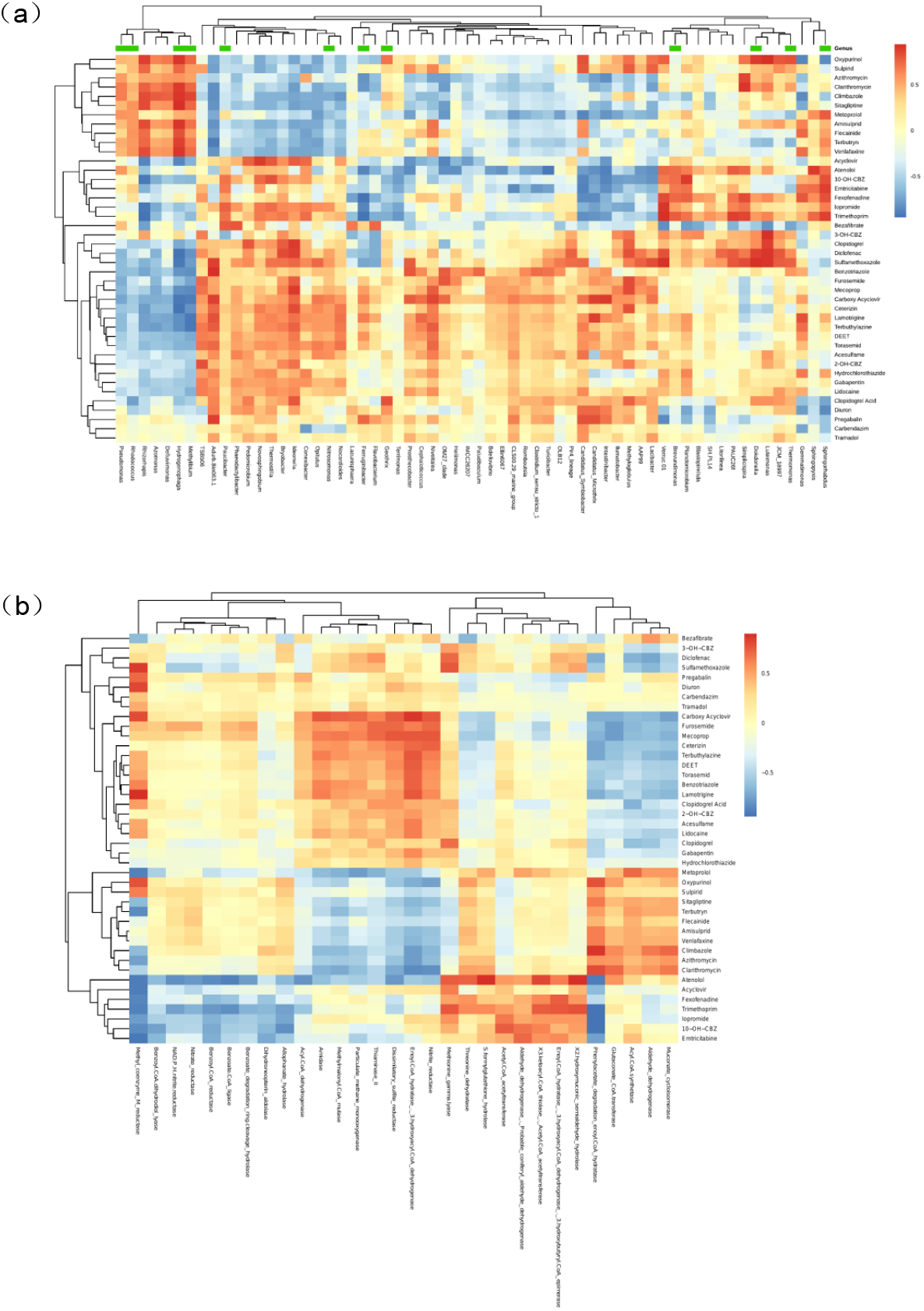
Heatmap showing correlations between biotransformation rate constants (*k_biol_*) of 45 TOrCs and (a) 62 genera; green bars represents the genera for which we obtained at least one MAG; (b) 30 significantly differential biotransformation related functions between the high biomass and the low biomass group. Cutoff is *p* < 0.05, abs(*r*) > 0.7, observation > 9. Average clustering was based on Euclidean distances.

By comparison with the integrated enzyme database of KEGG, we identified 20409 EC numbers from the metagenome sequences. Principle component analysis (PCA) based on the relative abundance of annotated functions demonstrated, similar to the taxonomic patterns described above, distinct clustering of different sand samples (Fig. 5). We can observe a separation between the HBG and LBG on axis 2 (PC2, 12%) with the exception of Friedrichshafen that shifted towards LBG, potentially caused by the low abundance of *Nitrospira*. PC1 (78%) distinguished two clusters of HBG, LBG and DW, which may due to the microbial structure differences between drinking water filters and wastewater filters. When comparing HBG with LBG, 1017 functions were over-represented in HBG and 1702 were over-represented in LBG, respectively (Fig. S5). From these, we selected those enzyme commission categories responsible for biocatalysis/biodegradation according to EAWAG-BBD database and analyzed their associations with TOrCs biotransformation. This resulted in 30 functions that showed a correlation with TOrCs biotransformation (with a cutoff of abs(*r*) > 0.7, adjusted *p* < 0.05, observation > 9) (Fig. 4b). Notably, enoyl-CoA hydratase (EC 4.2.1.17)/3-hydroxyacyl-CoA dehydrogenase (EC 1.1.1.35) encoding genes were positively correlated with eight compounds (i.e., benzotriazole, carboxy acyclovir, ceterizin, DEET, lamotrigine, mecoprop, terbuthylazine, and torasemid). Both, dissimilatory sulfite reductase (EC 1.8.99.3) and nitrite reductase (EC 1.7.2.1) genes, which encode for universal and essential enzymes in the sulfur and nitrogen cycle, were also correlated to the biotransformation of carboxy acyclovir, clarithromycin, furosemide and mecoprop. Further, a few functions showed a correlation to only a single compound removal rate. For example, amidase (EC 3.5.1.4) encoding genes were found to be only correlated to the *k_biol_* of carboxy acyclovir; threonine dehydratase (EC 4.3.1.19); acyl-CoA dehydrogenase, benzoate degradation ring-cleavage hydrolase, nitrate reductase (EC 1.7.2.1), S-formylglutathione hydrolase (EC 3.1.2.12) encoding genes were only correlated with the removal rate of atenolol.

**Figure 5.**
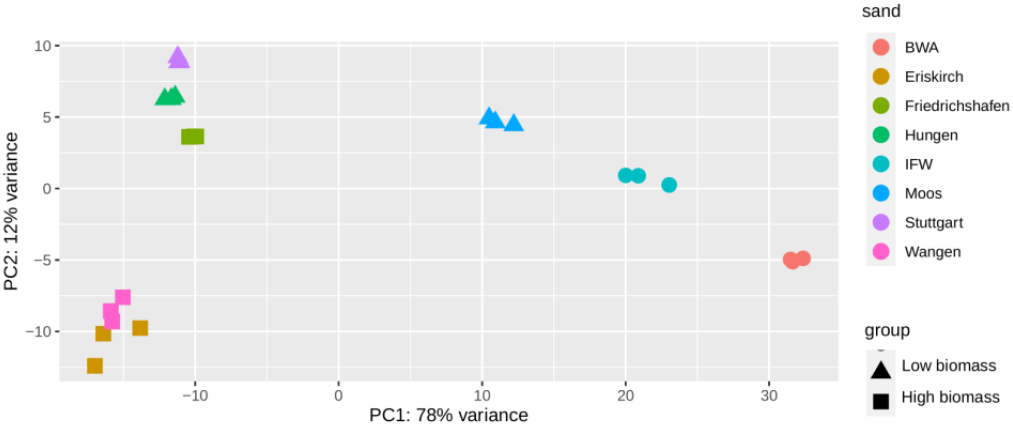
Principle component analysis (PCA) of eight sand filter materials based on the differential functions identified by DESeq2. Square symbol represents the high biomass group, triangle symbol represents the low biomass group, the remaining samples indicated by circles represent IFW and BWA.

### Biotransformation pathways analysis

Overall, there were 163 significantly differential abundant pathways annotated by HUMAnN2 comparing the HBG with LBG (see Fig. S6 for an overview excluding biosynthesis pathways). In the HBG, sulfate reduction, aromatic biogenic amine degradation and pathways regarding carbohydrate degradation (i.e. starch, glycogen, stachyose, D-galactose, galactose degradation) were overrepresented, the involved microorganisms were identified to be mainly Nitrospirae. In the LBG, energy metabolism pathways, such as the TCA cycle, Calvin-Benson-Bassham cycle, NAD/NADP-NADH/NADPH cytosolic interconversion, or the octane oxidation pathway were overrepresented.

We further inspected MAGs that taxonomically matched the genera for which we found correlations with individual *k_biol_* (see section above, Fig. 4a: genera with green bars) in order to see if these statistically as potential relevant genera contain biotransformation pathways. We annotated 37 MAGs in the KEGG Mapper for pathway reconstruction and we estimated the completeness of the pathways individually (Fig. 6). In the category of xenobiotics metabolism, MAGs classified to *Hydrogenophaga* showed the most complete pathways, especially in benzoate and furfural degradation. Steroid degradation pathway was prominent in *Sphingorhabdus* and *Pseudomonas*.

**Figure 6.**
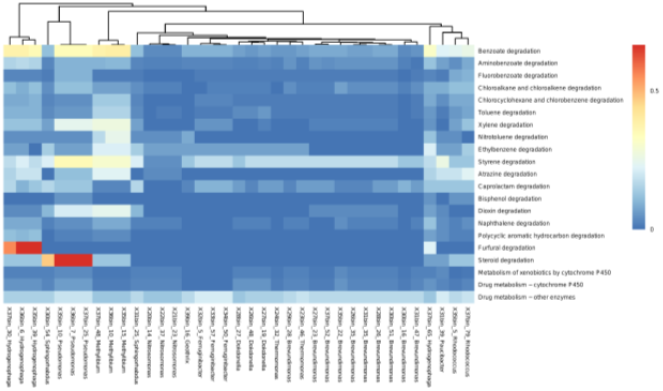
Heatmap showing the completeness of xenobiotics degradation pathways in 37 biotransfromation-correlated MAGs. Pathways are identified by “Reconstruct Pathway” tool in KEGG Mapper. Average clustering was based on Euclidean distances.

## Discussion

Currently, parameters such as redox conditions or biodegradable dissolved organic carbon, are used to evaluate the removal efficiencies of TOrCs (Bertelkamp *et al*. 2016; Torresi *et al*. 2019; Oberleitner *et al*. 2020), but new indicators directly associated to degradation processes (e.g., microorganisms, functional genes, transformation products) can be expected to be additional suitable tools for the prediction of the biotransformation potential and controlling removal performance. Here, we investigated the biotransformation of diverging TOrCs by eight biological active sand filter materials from wastewater and drinking water treatment plants, for which the metagenomic analysis of the microbial communities provided novel insights into the biological potential of TOrC transformations.

### Microbial biomass versus microbial community composition

We observed largely differing taxonomic and genetic profiles within each sand filter, accompanied by stark differences in the overall biotransformation performance (−1.4% vs 58% removal) across all investigated TOrCs. This may have led to the conclusion that functional microbial communities are the main predictor of TOrC transformations, however, most of these differences were eliminated after accounting for biomass, measured as ATP. Our results imply that the effect of biomass on the global TOrC transformation potential is foremost independent of the microbial community composition. We therefore had to partially reject our hypothesis that the microbial community composition is the major driver between global biotransformation potential in our experimental setup. However, on the other hand, the clear correlation of the compound matrix with the taxonomic matrix found by the Mantel test and later with single genera after normalization is evidence that although the global transformation potential is determined by biomass, the transformation of individual compounds is related to the taxonomic composition of the biological active sand filter system. After normalization, we could still identify significant differences for more than 15 different compounds (Table 1, e.g. clarithromycin, mecoprop, metoprolol, trimethoprim, furosemide, atenolol), which may be good candidates for a rather taxa or community specific degradation. Other studies already found indications that biomass is important for overall system performance, e.g. Liang *et al*. (2021) observed that the biomass increased with the running time of the reactors when also the TOrC removal increased, and Torresi *et al*. (2016) observed an enhanced TOrC removal with an increased biofilm thickness, although this feature was simultaneously attributed to increased diversity. In our study, however, alpha-diversity showed no such correlation (Table 2).

### Revision of biotransformation rate calculations

This indicates that we can predict the global transformation potential of (*ex situ*) materials from biofilters largely (explaining 76% to 84% of the *k_biol_* variation, or 90% of the variation for compound removal) by measuring the living biomass as ATP concentration. Further studies will be needed to assess if the relationship between ATP and *k_biol_* is linear or logarithmic (the two models were not significantly different (ANOVA, *p* = 0.24)). Biomass is used as a linear parameter for TOrCs biotransformation kinetics (see equation 1), and originally, the *k_biol_* was determined by the absolute abundance of the functional degraders of the respective TOrC (Bekins *et al*. 1998). In mixed microbial communities, however, this relationship may have to be reevaluated in the future.

When comparing frequently used biomass estimators for *k_biol_*, ATP, a biomass parameter that is frequently used in microbiology, has not been considered for *k_biol_* calculation of TOrC removal rates before. The biotransformation rate constants *k_biol_* were previously normalized by attached or suspended biomass as dry weight (Mazioti *et al*. 2015; Casas *et al*. 2015; Torresi *et al*. 2016), total suspended solids (Achermann *et al*. 2018), or DNA concentration (Liang *et al*. 2021). However, most methods are accompanied by certain biases. For example, dead and dormant cells will strongly influence DNA measurements and cell counts; or the biofilm EPS will bias DNA measurement (through external DNA), dry weight, and total carbon measurement. In our study, living and total cells (biased only by the different cell volumes) were still the best predictors after ATP (Table 2). Our findings suggest that at least for biofiltration systems the global potential for the biotransformation of TOrCs is more dependent on ATP than on other biomass indicators. Hence, ATP could have a profound impact when comparing biotransformation results across studies and treatments.

While this conclusion will hopefully change the way we analyze TOrCs biotransformations, ATP itself has been frequently used as an important biomass indicator in drinking water biofiltration systems for natural organic matter (NOM) transformation (Pharand *et al*. 2014; Chen *et al*. 2016; Kirisits *et al*. 2019). This is not surprising since NOM is used as a substrate by the microbial communities in various environments such as soil, sediment, marine, and freshwater (Kolehmainen *et al*. 2007; Huang *et al*. 2011; Diem *et al*. 2013; Simon *et al*. 2013). However, TOrCs are not considered as substrates per se and their concentrations are by definition generally considered as too low for biomass maintenance and growth and therefore efficient degradation. Co-metabolism through promiscuity or mixed substrate use can be a mechanism that lead to TOrC removal (Rauch-Williams *et al*. 2010, Hellauer *et al*. 2019).

### The potential role of rare taxa for biotransformations

In general, we divided the filters into three groups according to the clustering results, microbial community composition, and biomass estimates. The taxonomic composition between the HBG, LBG, DW, and also between single sand filters was different. HBG, though, shared Nitrospirae as highly abundant lineage (Fig. 3), a diverse and widespread group of often autotrophic, nitrite-oxidizing bacteria (Koch *et al*. 2015). Previous studies found that biotransformation of certain TOrCs can be related to ammonia oxidation activity of nitrifying activated sludge and biofilms in WWTPs (Helbling *et al*. 2012; Rattier *et al*. 2014; Men *et al*. 2017; Xu *et al*. 2017b). Moreover, asulam, carbendazim, fenhexamid, mianserin, and ranitidine showed biotransformation (16-85%) by the isolate *Nitrospira inopinata* (Han *et al*. 2019). Metagenome data of rapid gravity sand filter microorganisms also suggested that *Nitrospira* may serve as keystone species that drives the microbial ecosystems by providing organic carbon compounds and enable heterotrophic ammonium and carbon cycling (Palomo *et al*. 2016). In our system, we found no direct statistical linkage between *Nitrospira* and the biotransformation rate constants, however, it is possible that at least in the investigated biofilters Nitrospirae lineages as autotrophic species enables the establishment of a high microbial biomass, and thus enables the TOrC transformations through other microbial members. This also matches the observations by Liang *et al*. (2021), which indicates that the biotransformation may rely on rare community members of TOrCs-specific degraders, while the community of their moving bed reactor followed a progressive succession towards a Nitrospirae based climax community. The hypothesized positive and negative linkages of TOrC biotransformation with microbial genera pointed to rare biosphere microorganisms (< 2%), which exhibited most of the correlations (> 98%) with individual TOrCs removal rates. Although these correlations require experimental verification to test for causal relationships, these correlation-based hypotheses are in line with previous reports indicating that a small fraction of highly-specialized microorganisms, accounting for less than 0.1% of the microbial communities in biofilms, may be responsible for the TOrC transformation (Falås *et al*. 2018). One of the highly correlating genera was *Hydrogenophaga*, which showed a putative linkage to the removal of amisulprid, clarithromycin, climbazole, flecainide, oxypurinol, sitagliptine, sulpirid, terbutryn and venlafaxine (the top left cluster in Fig. 4a). This genus has also been previously reported to efficiently remove diclofenac, metoprolol, clarithromycin, erythromycin, atenolol and codeine (Kanaujiya *et al*. 2019).

In the drinking water sand filters DW, the biotransformation performance significantly differed from the wastewater filters. Here, *Pseudomonas* was found dominating DW, a common inhabitant of aquatic environments, including oligotrophic drinking water, lake water and surface water (Lopez *et al*. 2005; Huang *et al*. 2015; Nasreen *et al*. 2015). Although *Pseudomonas* has been reported to be able to degrade certain micropollutants (Li *et al*. 2010; Tezel *et al*. 2012; Devi *et al*. 2019), in our case, it also clustered in proximity to *Hydrogenophaga*, however, with more pronounced negative correlations with TOrCs.

### Enzymatic correlations with biotransformation rates

The relative abundance of functional genes and encoded biotransformation pathways also correlated with individual compounds and resulted in a clearer clustering than the microbial lineages. Enoyl-CoA hydratase/3-hydroxyacyl-CoA dehydrogenase was overrepresented in the HBG and was positively correlated with the *k_biol_* of eight TOrCs in our study. The enoyl-CoA hydratase is known to catalyze a β-oxidation substrate by adding hydroxyl groups and a proton to an unsaturated β-carbon of the molecule (Salgado *et al*. 2020). For instance, the degradation of ibuprofen can be initiated by enoyl-CoA hydratase with the introduction of hydroxyl groups, this enzyme was found to be up-regulated during the biodegradation process (Kjeldal *et al*. 2012). Enoyl-CoA hydratase was also identified in steroid estrogen degradation by bacterium *Serratia nematodiphila* (Zhao *et al*. 2020). Furthermore, Cameron *et al*. (2019) reported enoyl-CoA hydratase contributed to biofilm formation and the antibiotic tolerance, which also supported our findings that the high biomass promoted the TOrCs removal. Therefore, we suggest that a high abundance of enoyl-CoA hydratase/3-hydroxyacyl-CoA dehydrogenase might indicate a favorable functional potential for TOrCs biotransformation. Dissimilatory sulfite reductase (EC 1.8.99.3) and nitrite reductase (EC 1.7.2.1) were found to be correlated to the biotransformation of carboxy acyclovir, clarithromycin, furosemide and mecoprop in our study. In previous studies, sulfite reductase was reported to catalyze the cleavage of isoxazole and piperazinyl rings (Jia *et al*. 2019), and nitrite reductase was correlated with the biotransformation rate constant of some compounds like sulfamethoxazole, erythromycin and trimethoprim (Torresi *et al*. 2018). However, more studies are needed to further support this association.

## Limitations and perspectives

Our study was designed to compare the biological transformation potential of the filter materials *ex situ* under controlled conditions by applying laboratory scale transformation batch experiments. However, factors such as engineering design, redox potential, water retention times, feed/famine cycles that can influence the bioactivity, were explicitly not considered here. Moreover, we mainly focused on filter systems used for post-treatment of conventionally treated wastewater, and thus our results may be more representative for these types of rapid sand filters that have already experienced long-term exposure to TOrCs. The selection of 51 polar compounds maybe not fully representative of the full spectrum of TOrCs, but they cover a typical range of biodegradable and persistent compounds that can be found in wastewater (Ahmad *et al*. 2019), and the historical exposure of sand filters avoids artifacts due to falsely adapted or artificial assembled microbial communities. Since the selected TOrCs are relatively polar compounds and for many of them organic carbon normalized distribution coefficients have been reported to be rather low (Ternes *et al*. 2004; Stein *et al*. 2008; Ramil *et al*. 2010), sorption was considered negligible and we assumed that the observed removal was mainly attributed to transformation processes. Future studies should be conducted to answer the question whether ATP can also account for differences in *in situ* full-scale filters under natural retention times. Moreover, it should be validated if autotrophic taxa, such as Nitrospirae, can indeed act as primary carbon and energy deliverer that support potential rare indicator taxa such as *Hydrogenophaga* for individual TOrC transformations. Finally, our findings can be used as hypothesis for further looking into the details of TOrC biotransformation and its relationship with single taxa or whole communities.

## Conclusions

To summarize, we investigated the biotransformation performance of 51 TOrCs in eight sand filters from wastewater and drinking water treatment plants and established associations between the microorganisms, functional genes and the TOrCs biotransformation, which is also of high practical relevance as it could support the optimization and control of these systems. We conclude that

1. after normalization to microbial biomass, there was no significant difference of the average *k_biol_* between the main sand filter systems suggesting that biomass influences TOrC transformation globally;
2. the removal of individual compounds, however, was related to the taxonomic composition of the biological active sand filter system, indicated by individual *k_biol_* correlation with single genera, and by the global correlation of the microbial community composition with the normalized *k_biol_* matrix;
3. biotransformations of several TOrCs was rather correlated to rare biosphere lineages, e.g., *Hydrogenophaga* that had the most complete xenobiotics degradation pathways; on the enzymatic level, Enoyl-CoA hydratase/3-hydroxyacyl-CoA dehydrogenase, showed the broadest correlation with individual TOrC *k_biol_*; hence, these may be examples of suitable indicators for assessing biotransformation potentials;
4. the calculation of the *k_biol_* should be re-evaluated using a biomass marker for living cells (e.g. ATP) and traditional biomass estimators should not be used any more in transformation studies for normalization purposes.

## Data availability statement

All sequence data has been deposited at INSDC (with ENA: https://www.ebi.ac.uk/ena) under the accession number PRJEB43767. The ASV table, MAGs, and the DESeq2 output can be inspected in the supplementary material.

## Authors contributions

AW and DW designed and performed the experiment and the chemical analysis. LC, RL, and CW, analyzed the molecular data and made the statistical analysis. LC performed the metagenome assembly. CW and LC developed and wrote the initial draft of the manuscript and all authors improved the manuscript sequentially by several rounds of review.

## Acknowledgment

We would like to acknowledge the Leibniz-Rechenzentrum for providing computational support and Dr. Uwe Hübner for proof reading and a critical review of the manuscript. Dr. Claudia Beimfohr of Vermicon AG, Germany, is thanked for carrying out FISH analyses and cell count measurement.

