## Supplemental_material for "Microbial biomass, composition, and functions are responsible for the differential removal of trace organic chemicals in biofiltration systems"

**Supplementary material**

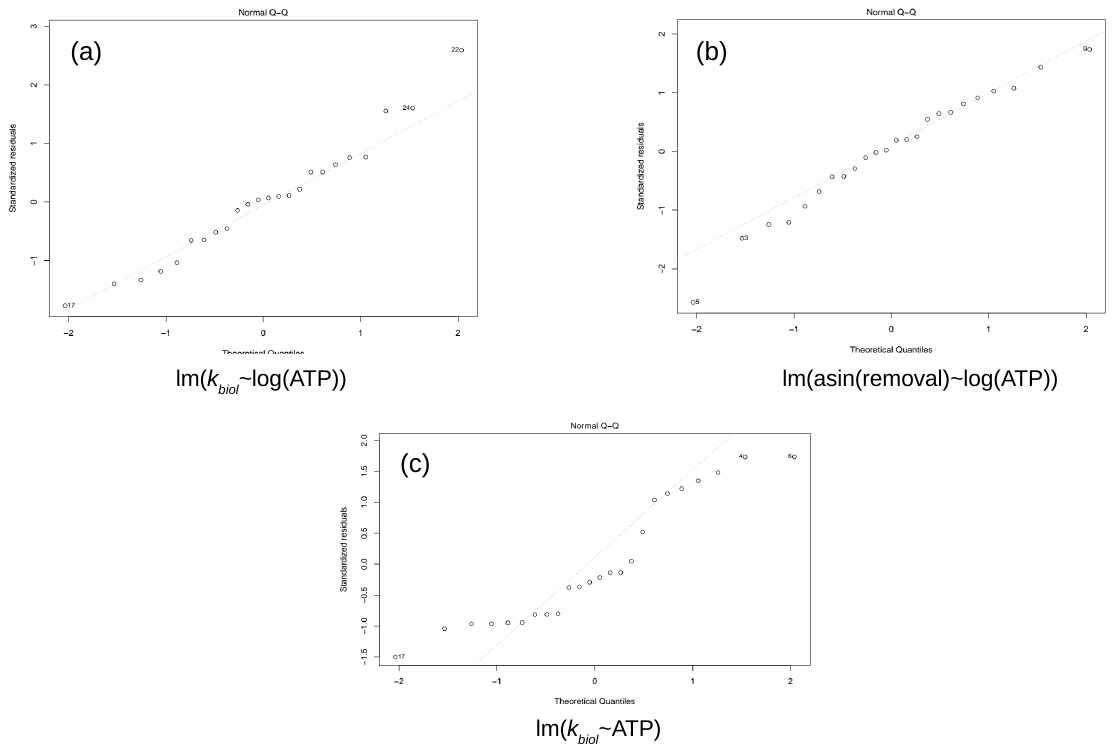

Figure S1. Normal QQ plot of residuals from linear model of (a) and (c) mean global k biol and ATP concentration (Fig. 1c); (b) mean global removal percentage and ATP (Fig. 1d).

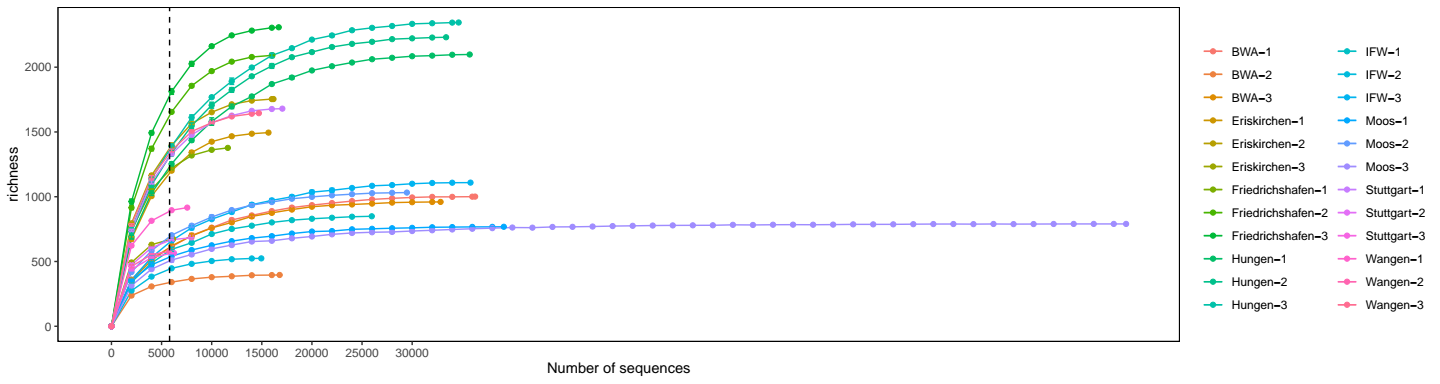

Figure S2. Rarefaction curve of 24 samples with pruning to 5799 reads. The x axis represents the number of sequences sampled while the y axis represents a measures of the species richness estimated with the Chao1 index.

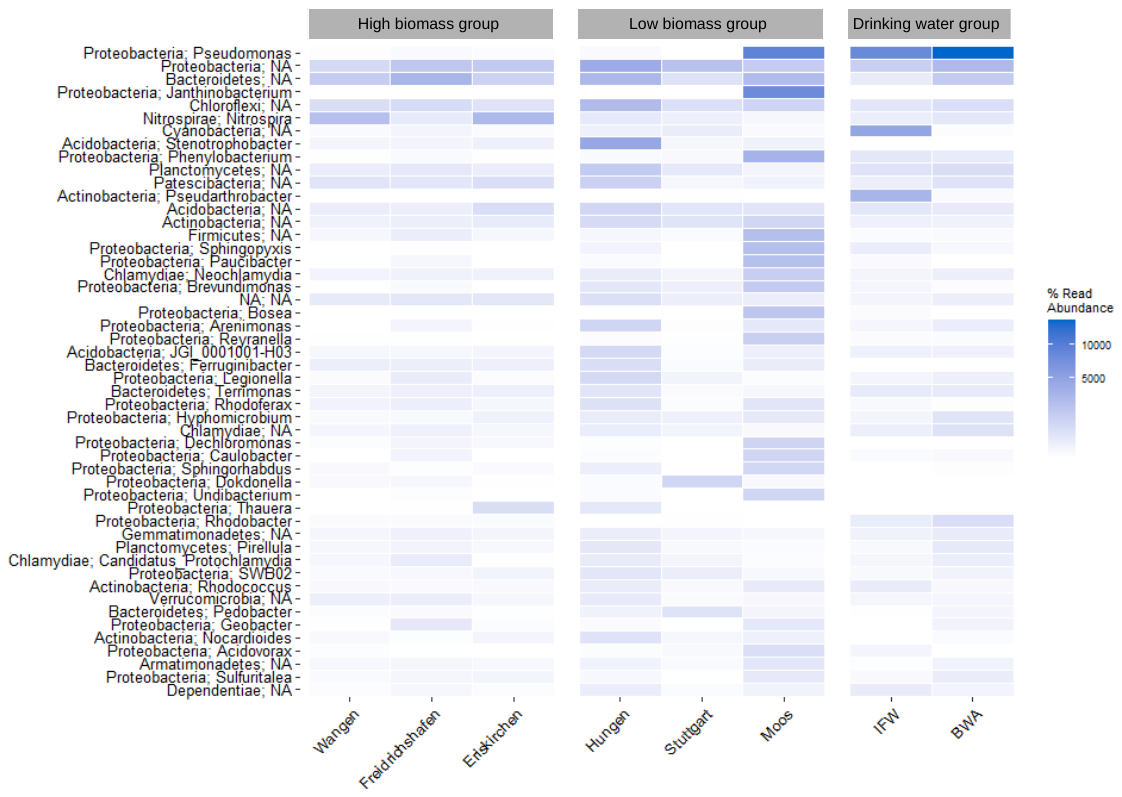

Figure S3. Taxonomic composition of eight sand filters at the phylum and genus level revealed by the most abundant 50 ASVs.

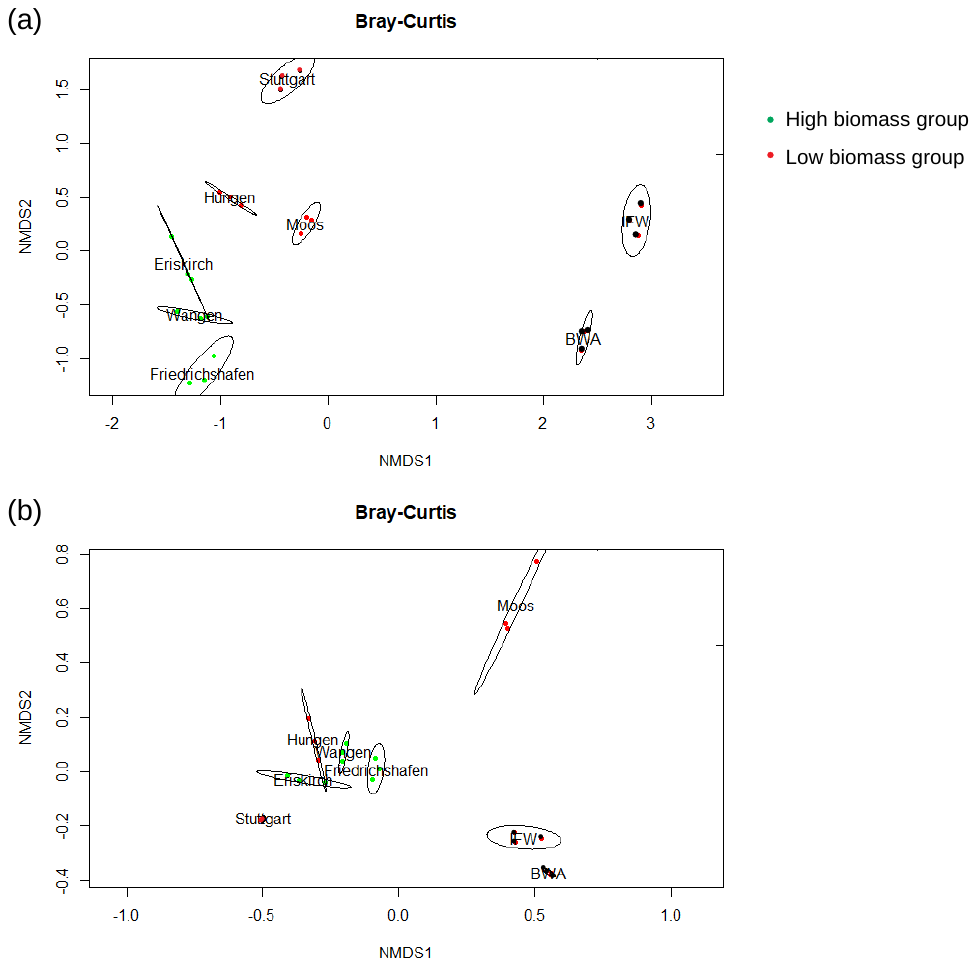

Figure S4. NMDS ordination based on (a) 16S rRNA data, (b) kWIP metagenome. Closer points imply more similar communities.

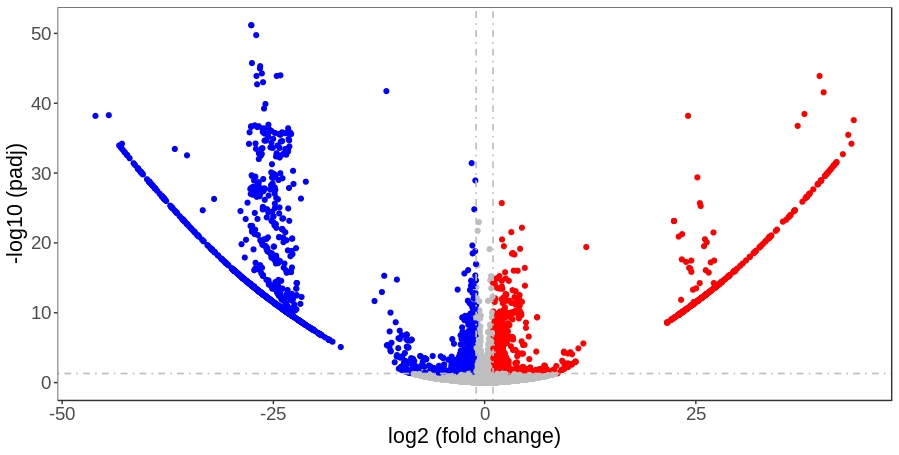

Figure S5. Volcano plot of significantly differential enzymes (high biomass group vs. low biomass group) annotated by SUPER-FOCUS. Red and blue color stand for overrepresented and downrepresented enzymes, respectively.

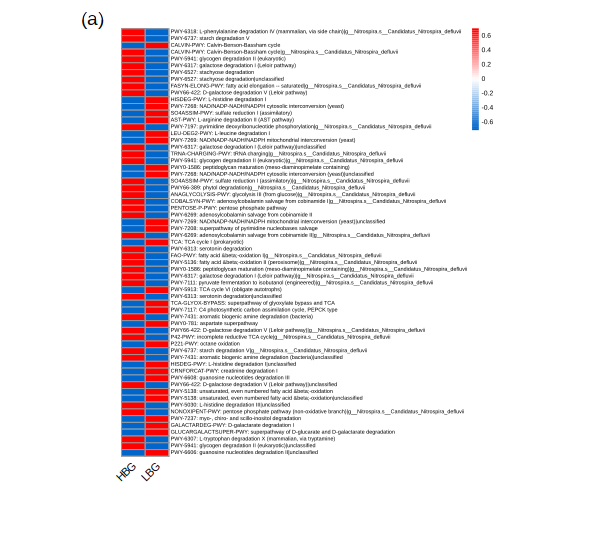

Figure S6. Significantly differential pathways (exclude biosynthesis) and involved microorganisms identified in the high biomass and low biomass group. The abundance of microbial pathways are profiled by HUMAnN2. The differential abundance analysis are performed by DESeq2.

Table S1. Number of reads passing through each step in DADA2 analysis.

| **Sample-ID** | **Sand** | **Input** | **Filtered** | **Denoised** | **Nonchim** |
| --- | --- | --- | --- | --- | --- |
| 17066-0148 |  | 82378 | 55100 | 47006 | 40443 |
| 17066-0149 | Eriskirchen | 80762 | 54972 | 45396 | 41461 |
| 17066-0150 |  | 38450 | 24443 | 18944 | 18024 |
| 17066-0151 |  | 42699 | 28196 | 22027 | 21150 |
| 17066-0152 | Wangen | 50510 | 34823 | 27461 | 26694 |
| 17066-0153 |  | 69300 | 49765 | 39628 | 37613 |
| 17066-0154 |  | 63962 | 46251 | 36894 | 34683 |
| 17066-0155 | Friedrichshafen | 77811 | 58515 | 46642 | 42394 |
| 17066-0156 |  | 80317 | 59070 | 47393 | 43990 |
| 17066-0157 |  | 80394 | 59661 | 50503 | 42278 |
| 17066-0158 | Stuttgart | 49646 | 33124 | 27570 | 22880 |
| 17066-0159 |  | 41278 | 26050 | 21532 | 19137 |
| 17066-0160 |  | 109682 | 84652 | 79077 | 60615 |
| 17066-0161 | Moos | 105174 | 81851 | 74387 | 57704 |
| 17066-0162 |  | 172658 | 137013 | 133166 | 96722 |
| 17066-0163 |  | 119764 | 92431 | 79352 | 65856 |
| 17066-0164 | Hungen | 126575 | 93532 | 79767 | 65891 |
| 17066-0165 |  | 129413 | 97233 | 82826 | 68928 |
| 17066-0166 |  | 103966 | 72224 | 65443 | 53531 |
| 17066-0167 | IFW | 58795 | 39965 | 35747 | 29746 |
| 17066-0168 |  | 122280 | 92120 | 83622 | 66135 |
| 17066-0169 |  | 129673 | 98212 | 90603 | 68554 |
| 17066-0170 | BWA | 65387 | 42892 | 39451 | 30652 |
| 17066-0171 |  | 105023 | 81551 | 74990 | 56908 |
